# Effect of *de novo* transcriptome assembly on transcript quantification

**DOI:** 10.1101/380998

**Authors:** Ping-Han Hsieh, Yen-Jen Oyang, Chien-Yu Chen

**Affiliations:** Graduate Institute of Biomedical Electronics and Bioinformatics, National Taiwan University, Taipei 10617, Taiwan; Department of Computer Science and Information Engineering, National Taiwan University, Taipei 10617, Taiwan; Department of Bio-Industrial Mechatronics Engineering, National Taiwan University, Taipei 10617, Taiwan; Genome and Systems Biology Program, National Taiwan University and Academia sinica, Taipei 10617, Taiwan

## Abstract

**Background:** Correct quantification of transcript expression is essential to understand the functional products of the genome in different physiological conditions and developmental stages. Recently, the development of high-throughput RNA sequencing (RNA-Seq) allows the researchers to perform transcriptome analysis for the organisms without the reference genome and transcriptome. For such projects, *de novo* transcriptome assembly must be carried out prior to quantification. However, a large number of erroneous contigs produced by the assemblers might result in unreliable estimation on the abundance of transcripts. In this regard, this study comprehensively investigates how assembly quality affects the performance of quantification for RNA-Seq analysis based on *de novo* transcriptome assembly.

**Results:** Several important factors that might seriously affect the accuracy of the RNA-Seq analysis were thoroughly discussed. First, we found that the assemblers perform comparatively well for the transcriptomes with lower biological complexity. Second, we examined the over-extended and incomplete contigs, and then demonstrated that assembly completeness has a strong impact on the estimation of contig abundance. Lastly, we investigated the behavior of the quantifiers with respect to sequence ambiguity which might be originally present in the transcriptome or accidentally produced by assemblers. The results suggest that the quantifiers often over-estimate the expression of family-collapse contigs and under-estimate the expression of duplicated contigs. For organisms without reference transcriptome, it remained challenging to detect the inaccurate abundance estimation on family-collapse contigs. On the contrary, we observed that the situation of under-estimation on duplicated contigs can be warned through analyzing the read distribution of the duplicated contigs.

**Conclusions:** In summary, we explicated the behavior of quantifiers when erroneous contigs are present and we outlined the potential problems that the assemblers might cause for the downstream analysis of RNA-Seq. We anticipate the analytic results conducted in this study provides valuable insights for future development of transcriptome assembly and quantification.

**Availability:** we proposed an open-source Python based package QuantEval that builds connected components for the assembled contigs based on sequence similarity and evaluates the quantification results for each connected component. The package can be downloaded from https://github.com/dn070017/QuantEval.

## Background

Quantification and comparison of transcript expression are essential to understanding the role of RNA in different physiological conditions or developmental stages. Such experiments and analyses are widely used in the studies of molecular biology. Over the past decades, several biological technologies have been developed to quantify the abundance of transcripts, such as expression microarray [1] and high-throughput RNA sequencing (RNA-Seq) [2]. For organisms with sufficient genomic information, the design of microarray provides a high throughput and cost-effective solution to examine transcript expression. On the other hand, RNA-Seq is superior in delivering lower background signals and larger dynamic ranges [3]. Despite the fact that many genome sequencing projects have been carried out, such as Genome10K [4], 5000 arthropod genomes initiative (i5K) [5] and Bird10K [6], whole genome studies are still demanding efforts for many research groups because of the immense cost and time-consuming process. For non-model organisms, the expression microarray needs to rely on cross-species hybridization [7]. On the contrary, RNA-Seq is more suitable owing to its capability of detecting novel transcripts without additional genomic information [3].

When the reference genome and transcriptome are not available, RNA-Seq reads are first used to reconstruct the transcriptome [8, 9]. Nowadays, many programs have been developed for *de novo* transcriptome assembly, such as Oases [10], rnaSPAdes [11], SOAPdenovo-Trans [12], Trans-ABySS [13] and Trinity [14]. After transcriptome sequences are reconstructed, quantification methods including BitSeq [15], Kallisto [16], RSEM [17] and Salmon [18] can be applied. These methods are able to inference the abundance of expression without the need of genomic sequences, using the number of RNA-Seq reads that overlap with the assembled contigs [9]. Nevertheless, quantification is much more challenging without reliable reference sequences because of the erroneous contigs produced by the assemblers, which often result from sequencing errors, insufficient sequencing depth and biological variability [19]. To address these problems, a great number of comparative studies have been published recently. While many studies evaluated transcriptome assembly [19-21] or quantification [22, 23] programs independently, few have discussed how transcriptome assembly influences the downstream quantification analysis. In 2013, Vijay, N., *et al*. performed an *in silico* assessment of RNA-Seq experiments. That study examined the impact of various aspects of sequencing reads on transcriptome assembly and differentially expressed genes (DEG) analysis [7], but the effect of redundant contigs and multiple-mapping reads on quantification was not well discussed. Another study conducted by Wang, S. and M. Gribskov evaluated the quality of assembled contigs and their effects on DEG analysis [24]. However, their study mainly focused on the evaluation of entire workflow from assembly, quantification, to DEG analysis, which makes it obscure to unravel how the erroneous contigs affect the authenticity of downstream analysis.

This study aims at comprehensively discussing the effect of biological complexity on transcriptome assembly, as well as the influence of incomplete and over-extended contigs on quantification. We used both *in silico* simulated and authentic RNA-Seq data from three species (yeast, dog, and mouse) to examine the effect of biological complexity on *de novo* transcriptome assembly. Three state-of-the-art assemblers, namely rnaSPAdes, Trinity and Trans-ABySS, were evaluated under different biological complexities based on TransRate [19] scores, which were previously proposed to assess the quality of *de novo* transcriptome assemblies using the alignments of sequencing reads to the assembled sequences. After *de novo* assembly, the reference transcripts were assigned to assembled contigs according to the BLASTn [25] alignments. Each transcript-contig alignment was then categorized based on accuracy, recovery and sequence ambiguity. Subsequently, we thoroughly examined the impact of erroneous contigs on the quantifiers Kallisto, RSEM and Salmon. By exploring the interplay between each stage in RNA-Seq analysis workflow, this study provides valuable insights into conducting RNA-Seq analysis and we anticipate these discoveries would be useful in the future development of assembly or quantification algorithms.

## Materials and Methods

### Datasets

Three experimental and three simulated RNA-Seq datasets were used in this study. Both experimental and simulated data included three species: yeast (*Saccharomyces cerevisiae*), dog (*Canis lupus familiaris*) and mouse (*Mus musculus*). The experimental datasets were collected from the Sequence Read Archive (SRA). The yeast dataset (SRR453566) was from the study of Nookaew *et al.* [26], comprising 5.5 million non-stranded paired-end reads cultivated under the batch condition. The dog dataset (SRR882109) was produced by Liu, *et al.* [27], comprising 20.8 million non-stranded paired-end reads sampled from normal mammary gland tissues of domestic dogs. Finally, the mouse dataset (SRR203276) was collected from the study of Grabherr, *et al* [14], containing 43.4 million stranded paired-end reads extracted from dendritic cells. For the simulated datasets, Flux Simulator (ver. 1.2.1) [28] was adopted to synthesize RNA reads for yeast, dog and mouse, respectively, based on the genomic sequences and annotations from the Ensembl database [29]. To facilitate the analysis, only the transcripts annotated as messenger RNA (mRNA) and with over 500 nucleotides in length were extracted.

The parameters used for the simulation are shown in Additional File 1: Table S1. In total, 81.7 million non-stranded paired-end reads were generated for the assembly and quantification analysis. The quality of both experimental and simulated datasets was examined using FastQC (ver. 0.11.5) [30] and the low-quality subsequences of the reads were trimmed using Trimmomatic (ver. 0.36) [31] with parameters *SLIDINGWINDOW:4:20 MINLEN:30*. The resultant RNA reads that were unable to maintain the paired relation were discarded. The detailed information of the processed RNA reads is provided in Additional File 1: Table S2 (The mean and standard deviation of the insert sizes were estimated based on the alignments produced by Burrows-Wheeler Aligner [32]).

### Expression Metrics

In order to evaluate the performance of transcript quantification, the ground truth of expression abundance for each transcript must be first determined. For simulated datasets, the numbers of the generated RNA reads for each transcript was recorded during the simulation process. Since transcriptome assemblers sometimes generate duplicated, incomplete or over-extended contigs, the metrics we use for quantifying expression must consider the normalization with respect to both sequence length and the number of total nucleotides. In this regard, the number of generated RNA reads was transformed into a simplified version of Transcripts per Million (TPM) [33] using the following equation:

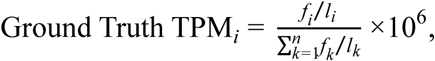

where *n* is the total number of transcripts, *f*_*i*_ is the number of RNA reads generated from transcript *i* and *l*_*i*_ is the effective length [34] of transcript *i*. In contrast, because the ground truth abundance for each RNA molecule is unknown for experimental datasets, we calculated the average TPM inferred by Kallisto (ver. 0.43.0), RSEM (ver. 1.2.31) (default parameters) and Salmon (ver. 0.8.2) for the reference transcript as the ground truth expression when evaluating the abundance of an assembled contig. Although the estimated expression might not perfectly reflect the real number of RNA molecules in a biological sample, it still provides valuable information when comparing the performance of quantification before and after *de novo* transcriptome assembly.

### *de novo* Transcriptome Assembly and Quantification

The processed RNA-Seq reads were assembled into contigs using the following three programs: (1) rnaSPAdes (ver. 3.11.1), (2) Trans-ABySS (ver. 1.5.5) along with ABySS (ver. 1.5.2) [35] and (3) Trinity (ver. 2.4.0) with default parameters. To minimize the effect of fragmented contigs, only the contigs with over 500 nucleotides in length were kept for the quantification analysis. The assemblies were evaluated based on the length of contigs, the number of recovered transcripts, the number of erroneous contigs and the evaluation scores provided by TransRate. The TransRate scores that we used in this study are the *score of bases covered score of good mapping score of not segmented* and *overall score*. The score of bases covered represents the proportion of nucleotide bases in a contig that are covered by reads. The score of good mapping represents the proportion of read pairs of which both reads are aligned in the correct orientation on a single contig. The score of not segmented represents the proportion of contigs that might be a chimera of multiple transcripts. Subsequently, the expression abundance for each contig was estimated using one alignment-based and two alignment-free quantifiers, namely (1) Bowtie2 (ver. 2.3.0) [36] (*--dpad 0 --gbar 99999999 --mp 1,1 --np 1 --score-min L,0,-0,1 -k 200 --sensitive --no-mixed --no-discordant*) followed by RSEM (ver. 1.2.31) (default parameter*s*); (2) Kallisto (ver. 0.43.0) (indexing with *-k 31* and quantifying with default parameters); and (3) Salmon (ver. 0.8.2) (indexing with *-k 31* and quantifying with default parameters).

### Transcript Assignment

For the purpose of comparing the estimated abundance of contigs with the ground truth expression from the corresponding transcripts, we assigned the reference transcripts to assembled contigs based on BLASTn (2.5.0+) [25] alignments. Here, only the high scoring pairs (HSPs) with identity over 70% and E-value smaller than 1E-5 were considered. We integrated the remained HSPs onto the coordinates of both transcript and contig to obtain the global alignment. Similar to a previous study [7], we calculated the *recovery* and *accuracy* for each global alignment, which refer to the proportion of matched nucleotides on the transcript and the proportion of correctly matched nucleotides on the contig respectively (Additional File 2: Fig. S1). Furthermore, we defined the overall *alignment score* as 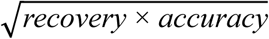. A transcript is assigned to a contig if either *accuracy* or *recovery* of the global alignment between them is above 90%. In this manner, we were able to identify all the corresponding transcripts for each contig. Note that it is possible that a contig can be associated with multiple transcripts, and a transcript can assign to multiple contigs as well. We considered multiple assignments here in order to understand the impact of redundant sequences on the quantification. Once the transcripts have been assigned to the contigs, we used the following equation to calculate the relative error of expression, in order to evaluate the quality of transcript quantification for each transcript-contig pair:

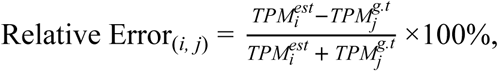

where the 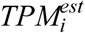 is the expression estimated by quantifiers for *contig*_*i*_, and the 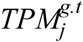 is the ground truth expression for *transcript*_*j*_ given (*contig*_*i*_, *transcript*_*j*_) is a valid global alignment (either *accuracy* or *recovery* of the global alignment between them is above 90%).

### Sequence Ambiguity

To determine the origin of the RNA-Seq reads that can be mapped to multiple transcripts is an important issue for the development of quantification algorithms. In this regard, it is of interest to understand the impact of sequence ambiguity on transcript quantification. We performed pairwise sequence alignment on both transcripts and contigs using BLASTn, respectively. Here, only the HSPs with identity over 70% and E-value smaller than 1E-5 were considered as potential ambiguity. In addition, to better explicate the relation between sequences that share similar subsequences, we build a connected component graph, where two sequences were grouped into the same connected component if the proportion of identical nucleotides between them is over 90% of the either sequence (Fig. 1). The size of a connected component is defined as the number of sequence members inside. We call the sequences in a connected component containing only one sequence as *unique sequence*. Furthermore, we used the read proportion of estimated abundance (RPEA) of a contig in a connected component to investigate the behavior of quantifier while ambiguous sequences are presented. Given *n* contigs *c*_*1*_, *c*_*2*_, *c*_*3*_*…c*_*n*_ in the same connected component *C*, the RPEA score for each contig is defined as follow:

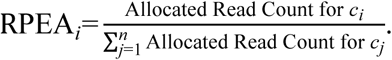

**Fig. 1:**
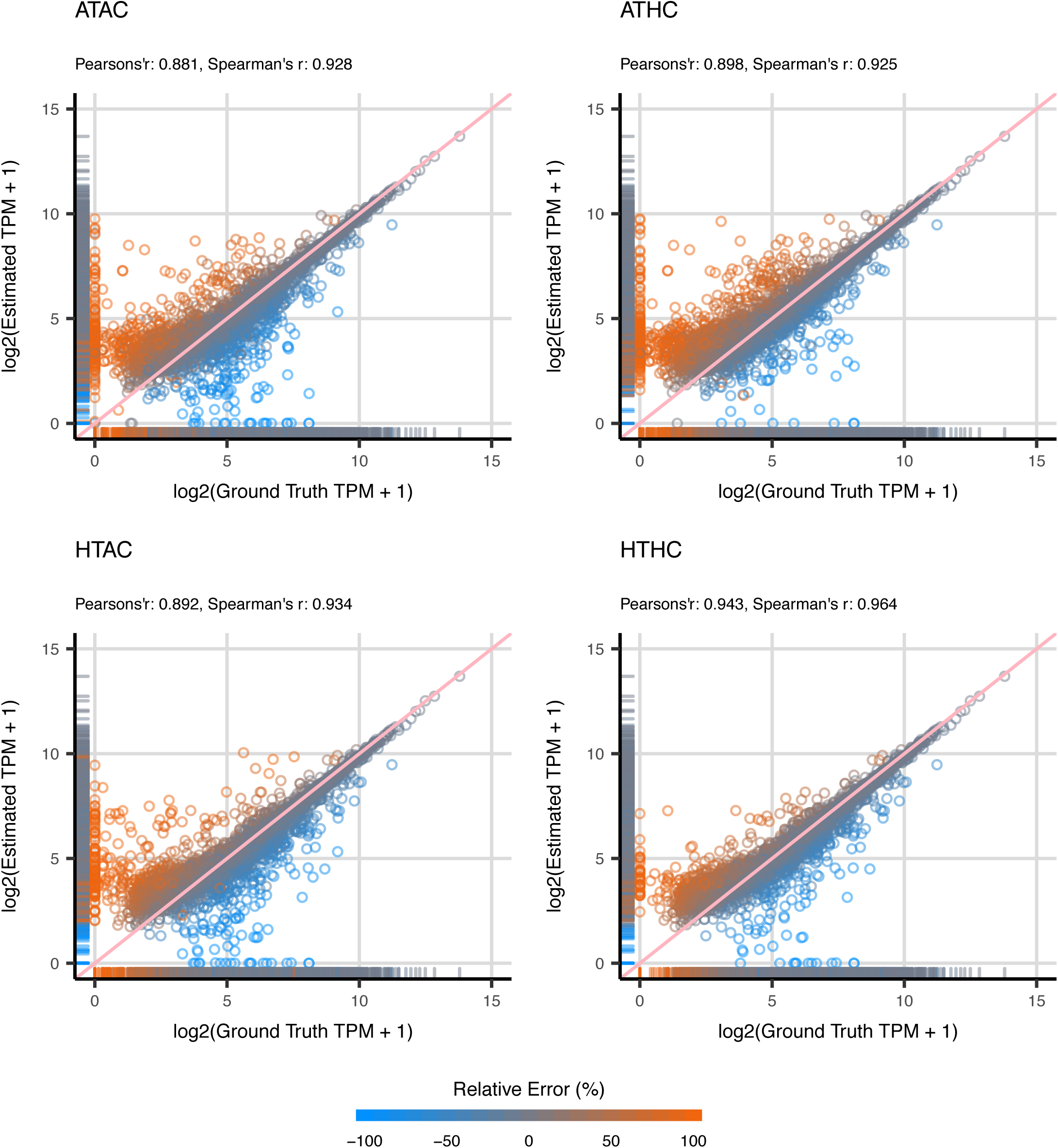
Construction of Ambiguity Network. The diagram illustrates how pairwise alignments in a contig set are employed to construct ambiguity networks. The ambiguity network is first initialized by given the contig sequences, creating a single cluster for each sequence. By analyzing the global alignments between contigs (the blue dot lines), the cluster in the network expands by joining two contig clusters at a time if the alignment length between the sequence is over 90% of the length for either sequence. In this study, the ambiguity network can be constructed for both contig and transcript sets. For the purpose of simplicity, we only illustrated the scenario for contigs in this figure.

If the highest RPEA in the connected component is close to 1, it suggests that the quantifier allocates all the reads in the connected component to one specific contig. In contrast, if the highest RPEA in the connected component is close to 1/*n*, then it suggests that the quantifier tends to allocate the reads evenly in the connected component.

### Contig Categories

The assembled contigs are categorized into five particular categories in this study: (1) *full-length*, (2) *incompleteness*, (3) *over-extension*, (4) *family-collapse* and (5) *duplication* (Fig. 2). The analysis of the first three categories were not affected by the factor of sequence ambiguity, allowing us to investigate the impact of assembly completeness on quantification independently. Given the length of contig *l*_*c*_ and the length of the corresponding transcript *l*_*t*_, the assembly completeness of a contig was examined through the difference in length:

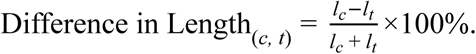

**Fig. 2:**
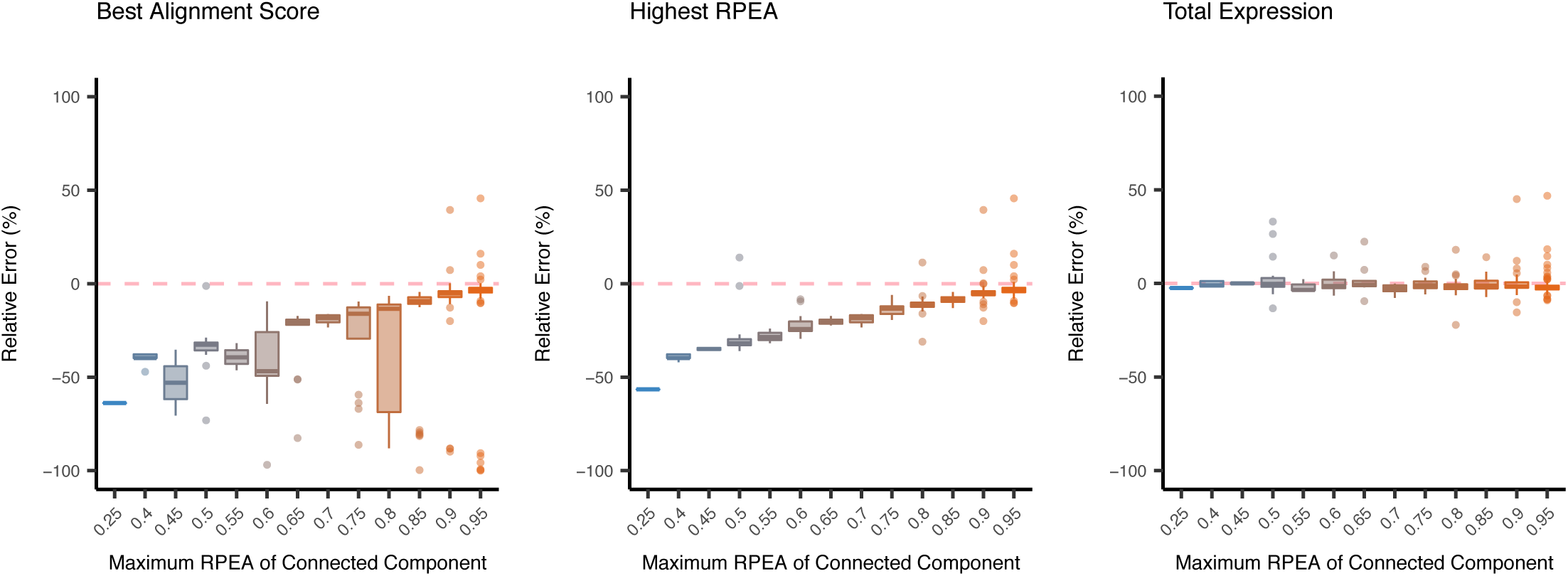
Examples of Contig Categories. The diagram gives examples for each contig category we analyzed throughout this study. The middle column shows an example for each category, while the right column portrays the relation of contigs and transcripts in network representation. The sequence nodes are connected together in solid line if they are in the same ambiguity cluster. On the other hand, a blue dot arrow represents the transcript assignment for the contigs. The analysis of contigs labeled as full-length, incompleteness and over-extended exclude the factor of sequence ambiguity. In contrast, family-collapse and duplication remove the potential effect of assembly completeness, focusing only on the impact of redundant or duplicated sequences.

In contrast, *family-collapse duplication* and *multiple-alignment* focused on the contigs that completely recovered the transcripts (*recovery* ≥ 90) but considered the influence of sequence ambiguity. To be more specific, *family-collapse* represents contigs which are assigned with multiple transcripts and *duplication* stands for the multiple contigs assigned by a single transcript. By examining these contigs, the problems caused by the assemblers that fail to distinguish similar transcripts from each other or generate a large number of redundant contigs are investigated. The detailed definitions for contig categories are provided in Additional File 1: Table S3.

## Results

### *de novo* Transcriptome Assembly

Based on pairwise BLASTn, yeast has the simplest transcriptome, with 94.11% of the transcripts sharing no similar subsequences with others. We call these sequences unique transcripts in the transcriptome. On the other hand, 66.29% of the dog transcripts are unique, while only 28.45% of the mouse transcripts are unique (Additional File 1: Table S4). First, we performed transcriptome assembly on the RNA reads of the three species with different biological complexities. We adopted three assemblers, rnaSPAdes, Trans-ABySS and Trinity, to construct contigs for both experimental and simulated RNA-Seq reads. Next, we built connected components for the assembled contigs. The statistics of sequence length and sequence ambiguity for the assembled contigs are shown in Additional File 1: Table S5, while the numbers of contigs that were categorized in each contig category are shown in Additional File 1: Table S6. The proportion of recovered transcripts (*recovery* ≥ 90) and accurate contigs (*accuracy* ≥ 90) is shown in Additional File 2: Fig. S2, and finally, the TransRate scores for each assembly are shown in Additional File 2: Fig. S3.

In general, rnaSPAdes constructed the least amount of contigs with the highest overall TransRate score for most of the datasets (5 among 6 datasets). As shown in Additional File 1: Table S5, rnaSPAdes also delivered the lowest average size of the connected components across species, suggesting that the contigs generated by rnaSPAdes are less redundant when compared to those from the other two assemblers. Trinity outperformed other assemblers in terms of N50 and the proportion of the recovered transcripts in all the simulated datasets. Despite the fact that Trinity generated longer contigs, the overall TransRate scores and the proportions of accurate contigs from Trinity assembly are marginally lower. Trans-ABySS constructed the contigs with comparatively high accuracy, with 66.37% of the contigs aligned with at least one transcript that show accuracy higher than 0.90. Nonetheless, Trans-ABySS constructed smaller numbers of unique and long contigs relatively. In other words, Trans-ABySS generated shorter contigs with more redundancy. The summary of the assemblies also demonstrates that the proportion of recovered transcripts are significantly higher in the datasets of yeast than that of dog or mouse. With higher complexity, it appears to become more difficult for the assemblers to properly reconstruct the transcriptome. For the estimated abundance of assembled contigs, the estimation made by quantifiers RSEM, Kallisto and Salmon shows considerably high consistency (Additional File 2: Fig. S4), with both Pearson’s and Spearman’s correlation coefficients higher than 0.95 between any of the two quantifiers.

### Impact of Assembly Completeness

In this study, the influence of *de novo* transcriptome assembly on expression quantification is mainly discussed with respect to two major issues: assembly completeness and sequence ambiguity. We would like to primarily look into the impact of assembly completeness on quantification in this section. In order to reduce the possible effect of sequence ambiguity generated by the assemblers, only the unique contigs (contigs in a connected component containing only itself) that are assigned with a single transcript were examined here. The unique contigs were further categorized into *full-length incompleteness* and *over-extension* (see Methods for detailed definitions). The reliability of quantification was examined based on the relative error for contigs with different extent of assembly completeness (Fig. 3).

**Fig. 3:**
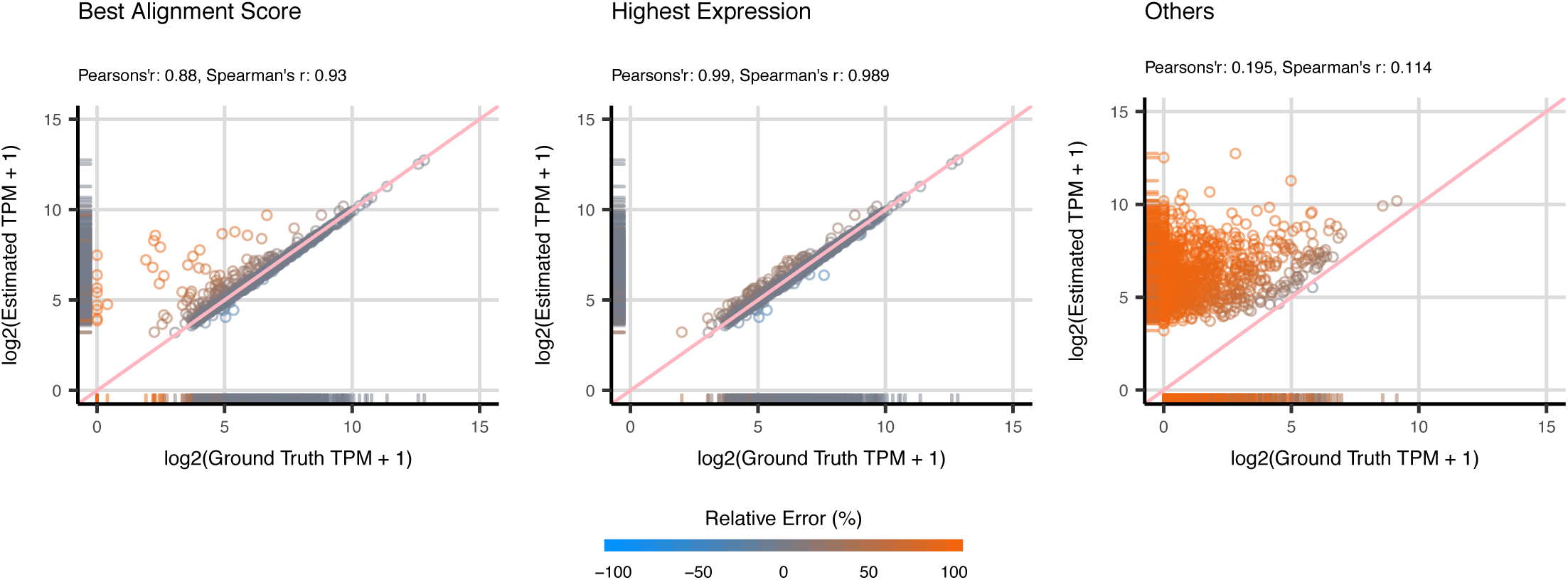
Quantification Errors for Unique Sequences. The box plots illustrate the relative quantification errors for unique contigs on the simulated datasets. The estimation of contig abundance is made by Kallisto based on Trinity assembly. The contigs are grouped by the extent of assembly completeness, and the numbers on the X-axis represent the lower bound of differences in length. For instance, the contigs located on −10 means that the percentage of difference in length is in the range of [-10, 0). The data is color-coded based on the contig categories. The box plots suggest that the estimation made on full-length contigs yield the smallest relative errors, while the incomplete contigs show over-estimation and over-extended contigs show under-estimation on quantification

In summary, full-length contigs show the lowest relative error of quantification, with the medians of error smaller than ±10%. The scatter plots and correlation coefficients also suggest that the estimated abundance of full-length contigs is highly reliable, with Pearson’s and Spearman’s correlation coefficient between the estimated and ground truth abundance both larger than 0.97 in all the datasets (Fig. 4, Fig. 5). The incomplete contigs yield slight over-estimation on the expression abundance. Overall, the quantification errors gradually increased as the assembly completeness decreased. This phenomenon can be observed more obviously on the dog and mouse datasets. When compared with the full-length contigs, the correlation coefficients are comparatively lower, ranging from 0.70 to 0.94 (Fig. 4, Fig. 5). Lastly, for the category of over-extended contigs, the quantifiers underestimated the expression abundance and the correlation coefficients slightly dropped (Fig. 4, Fig. 5). Nevertheless, the number of over-extended contigs is much fewer than those of other categories (Additional File 1: Table S6), which indicates that the assemblers did not overly extend the assembled contigs in most of the cases. In other words, only a limited number of contigs in the quantification will be affected in the practical RNA-Seq analysis. Although the TPM metrics has already been normalized for sequence length and total nucleotides, researchers might still need to be aware of the length bias caused by incomplete or over-extended contigs while using TPM as the metrics to estimate the expression of contigs.

**Fig. 4:**
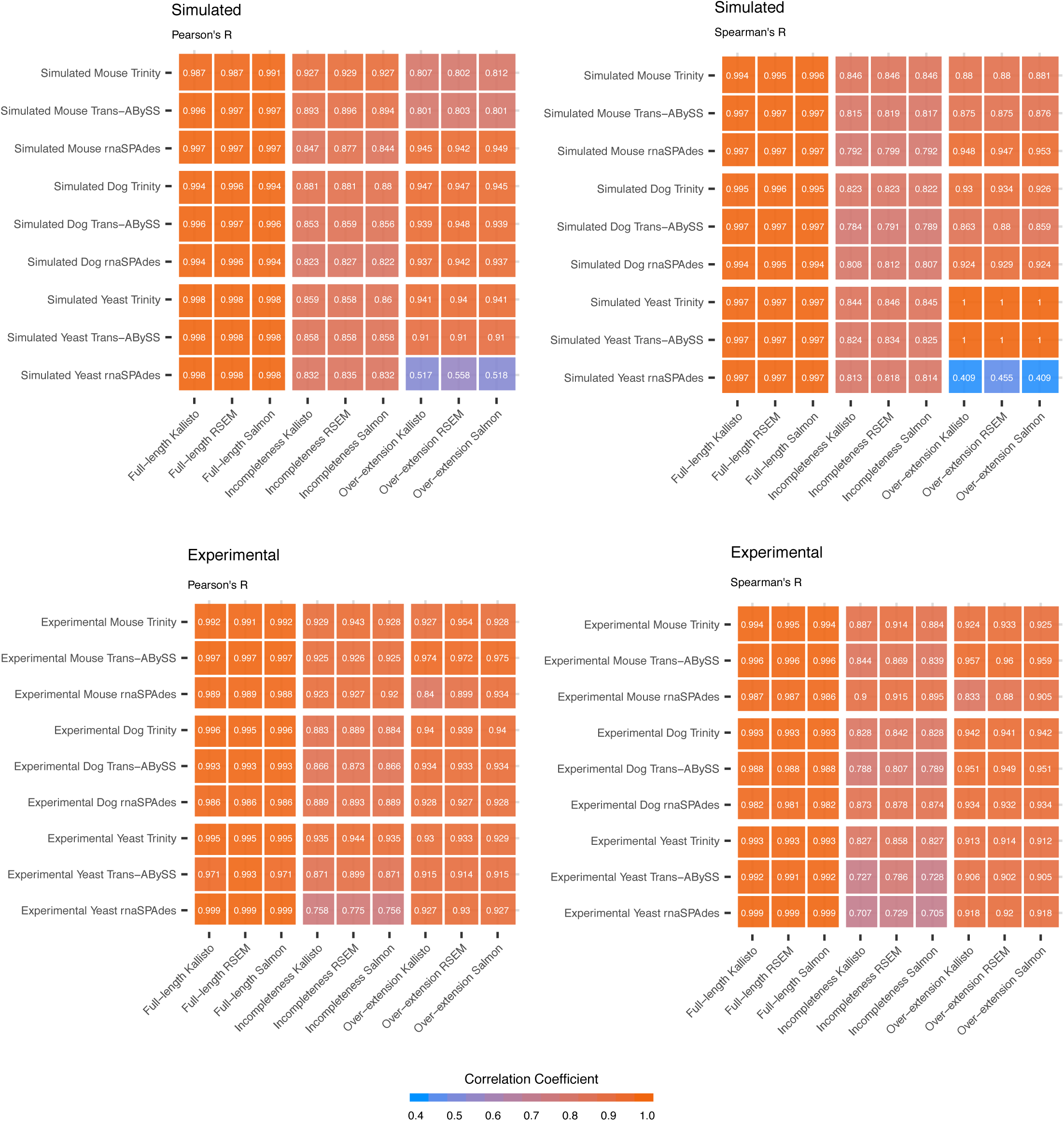
Scatter plots of Estimated Abundance and Ground Truth Expression for Unique Sequences. The scatter plots illustrate the estimated and ground truth abundance for contigs categorized as full-length, incompleteness and over-extension of the simulated dog datasets. The estimation of contig abundance is made by Kallisto based on Trinity assembly. The metrics are recorded in *log*_2_(*TPM* + 1). The data points are color-coded based on the relative quantification errors, with blue represents under-estimation and orange for over-estimation. In general, the estimation on expression for full-length contigs is highly reliable. There are some incomplete contigs with over-estimated abundance. Moreover, the correlation coefficients for the estimation of incomplete contigs are also relatively lower than that of full-length contigs. As for over-extended contigs, a marginal under-estimation in quantification can be observed.

**Fig. 5.**
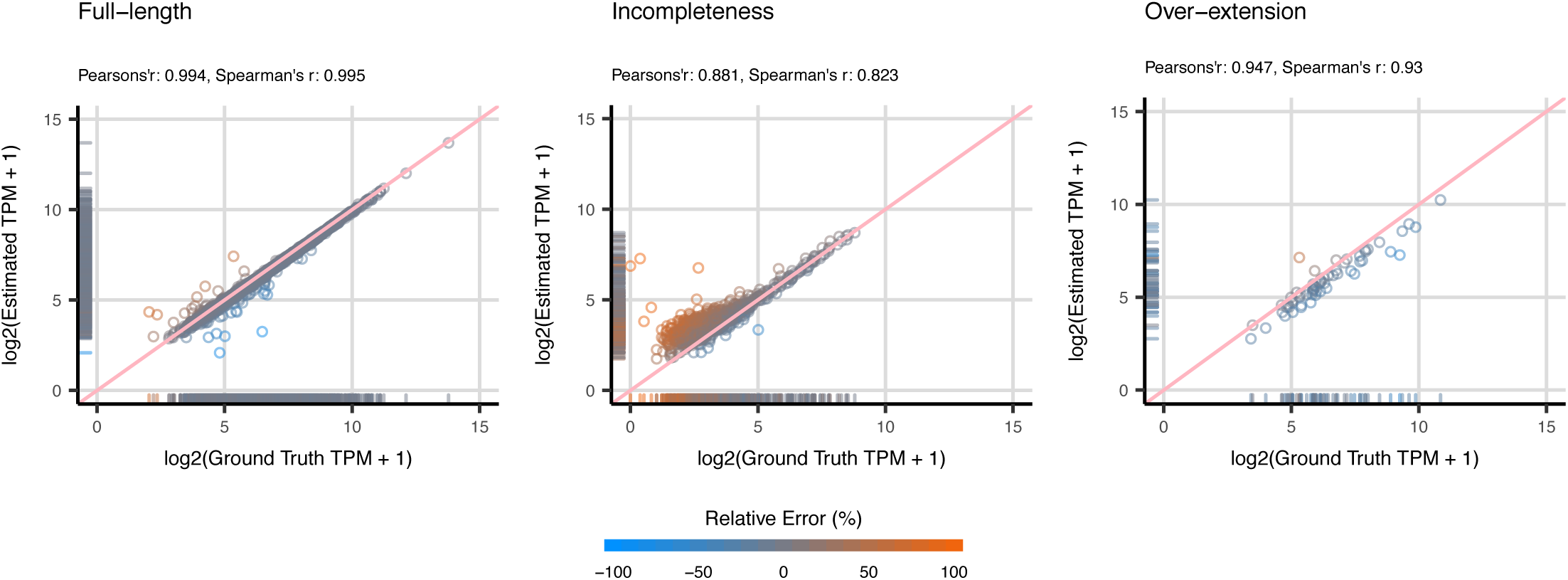
Correlation Coefficients between Estimated Abundance and Ground Truth Expression for Unique Sequences. The figures illustrate the Pearson’s and Spearman’s correlation coefficients between estimated abundance and ground truth expression. In general, the estimation based on full-length contigs have considerably high correlation with the ground truth expression of corresponding transcripts. In contrast, the incomplete and over-extended contigs show relatively lower correlation coefficients. There are significantly low correlation coefficients in the rnaSPAdes assembly based on simulated yeast data; however, due to a small number of data (n = 11), it should be careful to draw such conclusion based on this dataset.

### Impact of Sequence Ambiguity

In this section, the impact of sequence ambiguity on quantification was thoroughly discussed. Similarly, to reduce the compound effect from assembly completeness, only the accurately assembled contigs (*accuracy* ≥ 90) were examined here. In the first part of this sub-section, we looked through the reliability of quantification when the assemblers report only one contig for many similar transcripts, denoted as *family-collapse*. In the second part, we examined the impact of contigs with similar sequence content which are assigned with the same transcript, denoted as *duplication* (see Methods for detailed definition). By using these contigs, we examined the behavior of the quantification algorithms while sequence ambiguity is present in the assembly.

For the contigs categorized as family-collapse, it is much more difficult to analyze the accuracy of quantification because of multiple transcripts being assigned to a contig. Based on our observation, there are in average 2 to 3.16 transcripts being assigned to a family-collapse contig across the six datasets. Since there is only one contig that were assigned by multiple transcripts, we would like to find out of which transcript expression delivers the estimated abundance of the contig actually reflects. To our surprise, the estimated abundance is closer to the transcript with the highest expression rather than the one with highest alignment score (Fig. 6, Additional File 2: Fig. S5).

**Fig. 6:**
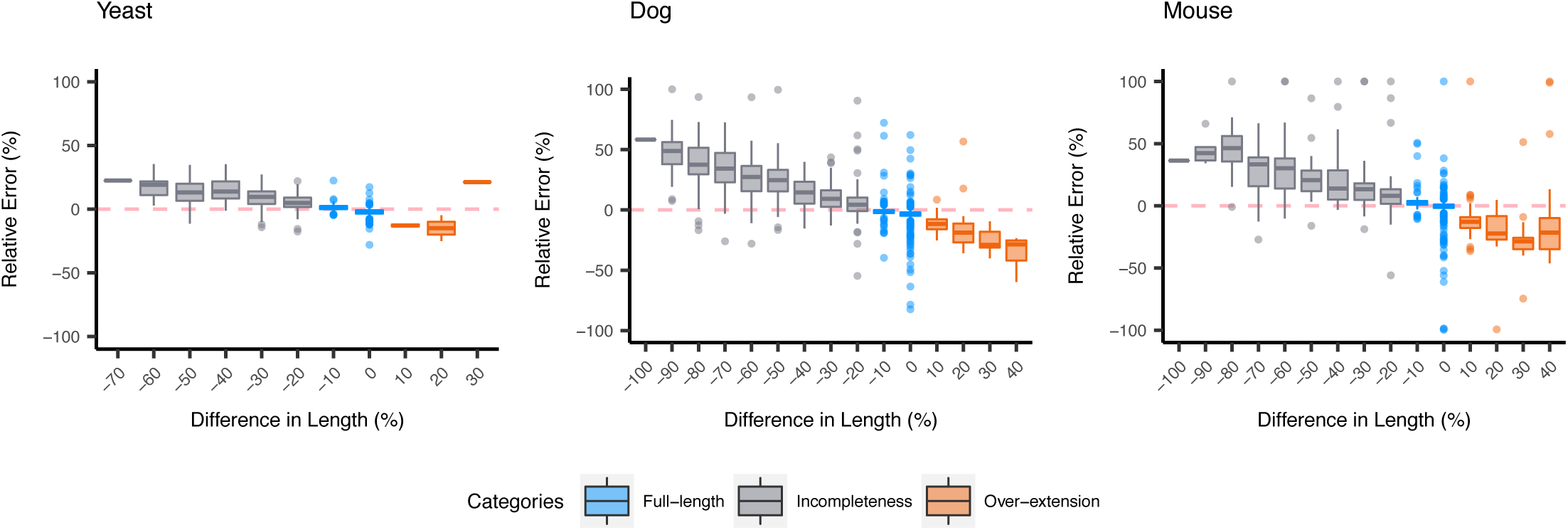
Scatter plots of Estimated Abundance and Ground Truth Expression for Family-Collapse Sequences. The scatter plots illustrate the estimated and ground truth abundance for contigs categorized as family-collapse of the simulated dog dataset. The estimation of contig abundance is made by Kallisto based on Trinity assembly. The metrics are recorded in *log*_2_(*TPM* + 1). The data points are color-coded based on the relative quantification errors, with blue represents under-estimation and orange for over-estimation. Since there are more than one transcripts correspond to one contig, we categorized the expression of corresponding transcript into (1) transcript with the maximum alignment score with respect to the contig, (2) transcript with the highest expression in the family, and (3) others. In general, the estimated abundance of the contig actually reflect to the transcript with the highest expression.

In contrast with family-collapse, duplication represents the redundant contigs which are clustered into the same connected component and are assigned with a single transcript. Here, we use the maximum estimated abundance in the connected component to investigate the behavior of quantifiers (see Methods for detailed definition). We observed that quantification algorithms tend to allocate most of the RNA reads to a single contig within the connected component in most of the cases (15 among 18 of the datasets show that over 50% of the connected component has one contig with the proportion of estimated abundance over 75%) (Additional File 2: Fig. S6). Furthermore, we would like to understand which estimated abundance of the contig in the connected component can accurately reflect the expression of corresponding transcript. Here, we used three approach to select the estimated abundance of the contig in the connected component: (1) the contigs with the highest alignment score, (2) the contigs with the highest RPEA and (3) the total expression of connected component of the contigs. Consequently, we found that the estimated abundance of contigs that were allocated with the most amount of RNA reads in the connected component show significantly low quantification error with the transcript expression. However, in the cases when the quantifiers distribute the RNA reads evenly to the duplicated contigs, the ground truth expression for the transcripts cannot be accurately represented by the estimated abundance of contigs. (Fig. 7). To address this problem, it is advisable to use the total expression of the connected component for duplicated contigs to measure the expression of corresponding transcripts.

**Fig. 7:**
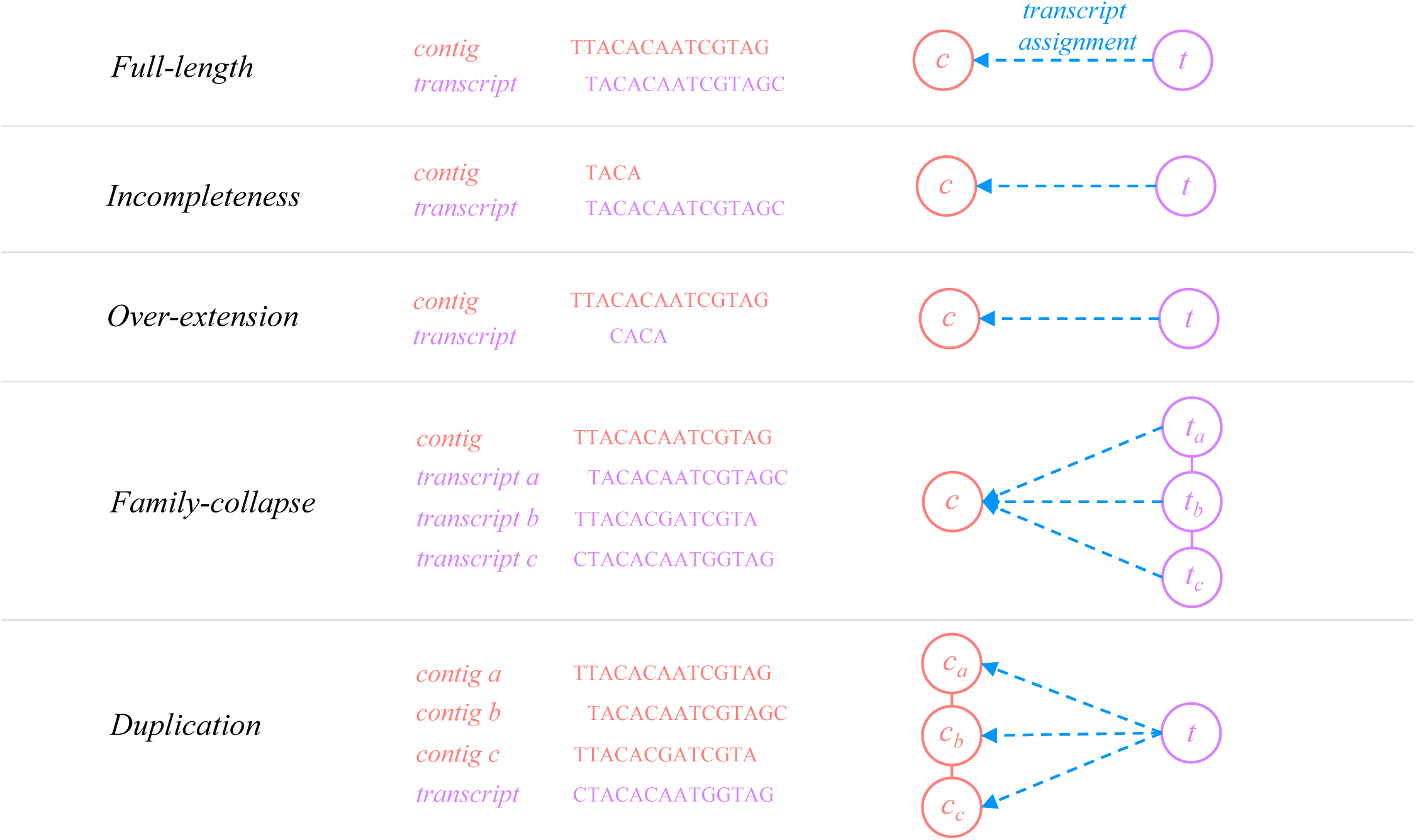
Box Plots for the Relative Errors of Duplicated Contigs. The box plots illustrate the relative quantification errors for duplicated contigs of the experimental mouse dataset. The contigs are grouped by the maximum RPEA in the connected component, and the numbers on the X-axis represent the lower bound of the proportion. For instance, the contigs located on 0.45 means that the maximum RPEA of the connected component is in the range of [0.45, 0.50). Since there are more than one contigs that are assigned by the same transcript, we would like to find out which contigs’ estimated abundance can accurately reflect the expression of the transcript. Here, we categorized the quantification errors into three categories: (1) transcript is assigned to the contig with the highest alignment score, (2) transcript is assigned to the contig that are allocated with the most RNA reads and (3) transcript expression adopts the total expression of the connected component of the associated contigs. The box plots suggest that contig with the highest alignment score or the highest estimation made within the connected component have considerably lower quantification errors if most of the reads are assigned to one specific contig (higher maximum RPEA). However, when the quantifiers allocate the RNA reads evenly to the contigs within the connected component, it is advisable to use the total expression of the connected component instead in order to get the accurate estimation for the expression of transcripts.

## Discussions

### Component-level Quantification

Based on the observation discussed in the previous section, we first discovered that once the assemblers failed to distinguish the RNA reads generated from similar transcripts and reported a single merged contig, the estimated abundance of family-collapse contig often reflect the expression of the transcript generating the most amount of RNA reads. In contrast, to estimate the expression of the transcript associated with duplicated contigs, researchers are suggested to use the abundance of the contig that is allocated with the majority of the RNA-reads (highest RPEA score) in order to get accurate estimation. Nevertheless, in most of the practical RNA-Seq analysis, the information of the transcriptome sequence is not available for non-model organisms. Therefore, it is challenging to detect whether family-collapse contigs or duplicated contigs emerge when performing contig annotation (transcript assignment) after assembly. Here, we looked through all the potential assignment of transcripts for each contig and we demonstrated four different strategies: (1) selecting the transcripts with the highest alignment score for each contig, followed by selecting the contig with the highest alignment score for each transcript (ATAC), (2) selecting the transcripts with the highest alignment score for each contig, followed by selecting the contig with the highest RPEA score in the connected component (ATHC), (3) selecting the transcripts with the highest expression in the connected component, followed by selecting the contig with the highest alignment score for each transcript (HTAC), and (4) selecting the transcripts with the highest expression in the connected component, followed by selecting the contig with the highest RPEA score in the connected component (HTHC). As a result, it seems to be more appropriate to use the estimated abundance of contigs with the highest RPEA score in the connected component to reflect the expression of transcripts (Fig. 8). Despite the fact that this strategy abandons quantifying individual transcripts, many advantages emerge such as higher accuracy on the quantification, reliable read inference, robust statistical performance and clear interpretation of the data [37]. More importantly, this strategy does not have to deal with the problem resulting from multiple transcript assignment in contig annotation.

**Fig. 8:**
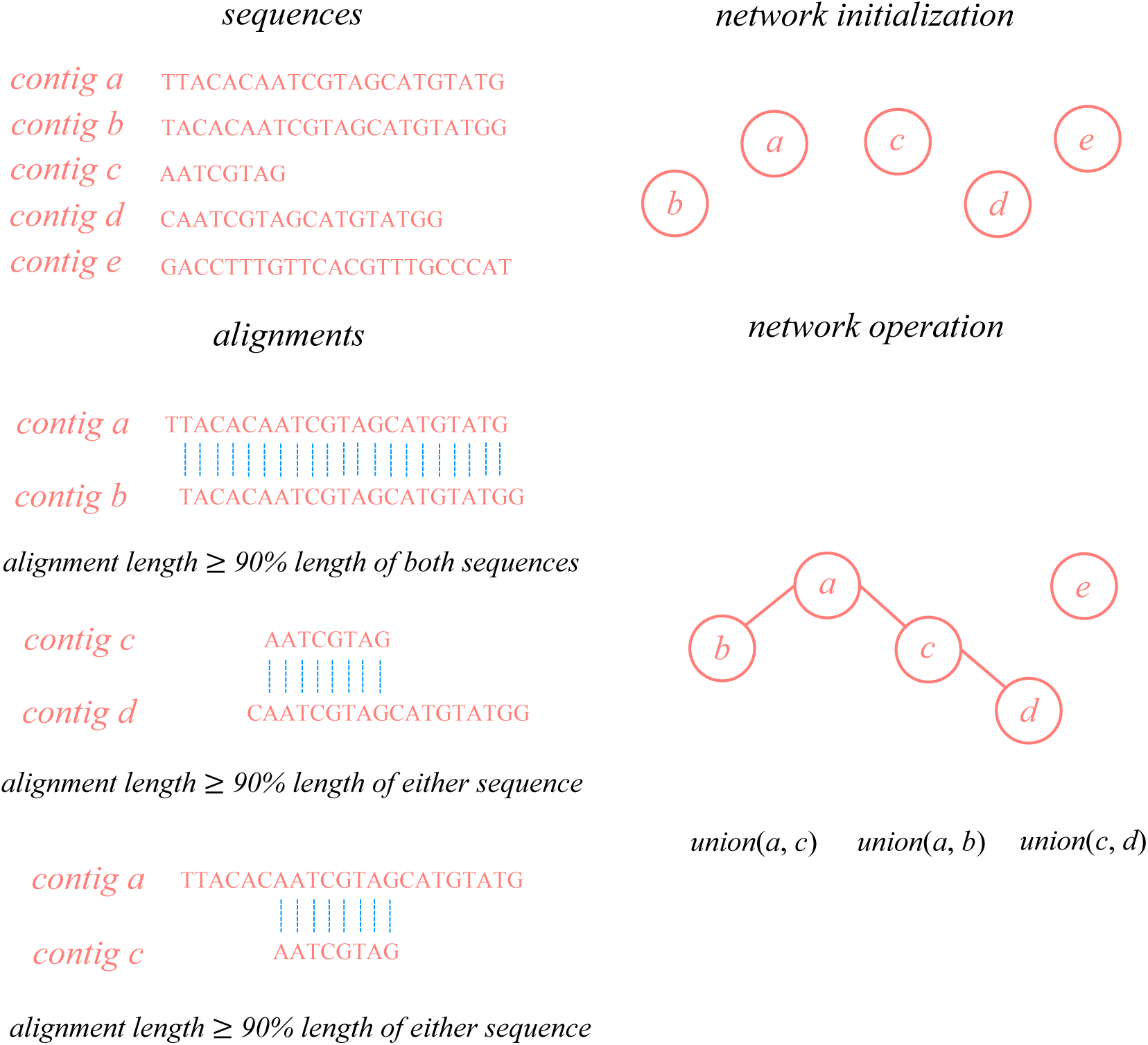
Scatter plots of Estimated Abundance and Ground Truth Expression for All Sequences. The scatter plots illustrate the estimated and ground truth abundance for all the contigs of the simulated dog dataset. The estimation of contig abundance is made by Kallisto based on Trinity assembly. The metrics are recorded in *log*_2_(*TPM* + 1). The data points are color-coded based on the relative quantification errors, with blue represents under-estimation and orange for over-estimation. We looked through all the valid assignment of transcripts for each contig and we demonstrated the correlation for four different strategies: (1) selecting the transcripts with the highest alignment score for each contig, followed by selecting the contig with the highest alignment score for each transcript (ATAC), (2) selecting the transcripts with the highest alignment score for each contig, followed by selecting the contig with the highest RPEA score in the connected component (ATHC), (3) selecting the transcripts with the highest expression in the connected component, followed by selecting the contig with the highest alignment score for each transcript (HTAC), and (4) selecting the transcripts with the highest expression in the connected component, followed by selecting the contig with the highest RPEA score in the connected component (HTHC) We would like to understand whether the estimated abundance for both methods can be used to accurately estimate the expression of transcripts. In general, the estimated abundance of the contig with the highest number of allocated reads in the connected component (HTHC) yield a more accurate estimation.

### Selection of Programs

We argue that the difficulties discussed in this research emerge mainly because the goals for each step of the analysis is not specifically designed to suit *de novo* RNA-Seq analysis. Most of the assemblers are optimized to reconstruct the whole transcriptome, which sometimes leads to many false predictions on SNPs or isoforms. These artificial or incorrect contigs thereafter deteriorate the accuracy of quantifiers because most of the quantification algorithms infer the expression based on the overlapping relation between RNA-Seq reads and the given sequences (transcripts or contigs). Furthermore, the annotation step is performed based on sequence alignment without considering the abundance of expression. This observation demonstrates that the selection of the programs has a strong impact on the *de novo* RNA-Seq analysis. For instance, to minimize the effect of assembly completeness on quantification, assemblers that construct the sequences with appropriate length are preferable. On the other hand, to reduce the problem in contig annotation resulting from sequence ambiguity, the assemblers that report fewer false prediction for SNPs or isoforms are better choices. The performance for the quantifiers we analyzed throughout this study demonstrates high consistency in terms of accuracy of quantification. Therefore, we recommended to use the alignment independent quantifiers such as Kallisto and Salmon for significantly lower computational time [16, 18]. The selection of programs for contig annotation is relatively irrelevant because the bottleneck mainly lies in the insufficient transcript information. Since there are no reference transcripts available in the practical *de novo* RNA-Seq analysis, most of the research utilizes the protein sequences from closely related species because of comparatively higher conservation in protein sequences. However, to align the contigs with proteins of other species might reduce the precision and accuracy of sequence alignment, which makes it more difficult to find the correct annotation for the assembled contigs.

### Other Important Factors in RNA-Seq Analysis

Although we mainly discussed the impact of assembly completeness and sequence ambiguity on quantification and annotation throughout this study, there are many other factors that might as well result in unreliable experimental results in the practical *de novo* RNA-Seq analysis. For instance, the read length or the fragment size directly determine the maximum length of the nucleotides that overlap with the reference sequences for each read pair. Therefore, the number of RNA-Seq reads that can be aligned to multiple origins of the transcripts reduces when longer reads are adopted, which mitigates the problem of sequence ambiguity on the inference of the origin of the reads [33, 34]. The strand specificity provides the information for the strand of the RNA-Seq reads, which improves the precision of sequence alignment and quantification [38]. Last but not least, the sequence-specific and positional bias derived from library construction might lead to RNA-Seq reads that over- or under-represent the number of transcripts in the molecules In this regard, it is important to model the fragment bias in the process of quantification [39]. However, considering all the combined effects of the factors mentioned above will significantly increases the complexity of this research. Therefore, we generated non-stranded RNA-Seq with uniform read length, and the fragment bias for the reads were excluded. Moreover, the start coordinates of the RNA-Seq reads uniformly distributed. By this means, we excluded the potential impact of these factors and focused only on the effect of assembly completeness and sequence ambiguity. But the importance of these factors should not be ignored in the practical *de novo* RNA-Seq analysis.

### Sequencing Technologies

In this section, we briefly highlight two sequencing technologies that provide another new perspective and mitigate the problems in RNA-Seq analysis: single-cell RNA sequencing (scRNA-Seq) and long-read RNA sequencing. By isolating individual cells prior to sequencing, scRNA-Seq technology allows the researchers to extract mRNA from a single cell. The development of scRNA-Seq allows the researchers to identify uncharacterized cell types, to observe the stochastic nature of gene expression, and to compare the expression profiles for the individual cells within the population [9, 40]. Nevertheless, the number of reads generated using scRNA-Seq is relatively small since only a small number of mRNA molecules are expressed in a single cell. This drawback makes it difficult to reconstruct the transcriptome for each cell [40]. Moreover, assembly algorithms that take the advantages of unique molecular identifiers (UMIs) for each read are still in demand. In the *de novo* RNA-Seq analysis, it is advisable to aggregate the scRNA-Seq from all cells as the bulk RNA-Seq reads to reconstruct the transcriptome and subsequently estimate the abundance of each cell subsequently. The restriction makes the exploitation of scRNA-Seq mainly limit to species with highly-curated transcript sequences available.

Another technology that developed with increasing interests recently is long-read RNA sequencing. Unlike next-generation RNA-Seq technology, long-read RNA sequencing generates a single read for each mRNA molecule in real-time, which results in considerably longer RNA-Seq data that allows the full-length reconstruction for transcripts without the need of assembly [9]. Although the demanding cost for sequencing in higher coverage makes it hard to be considered for quantification at this moment, this technology still provides an extraordinary breakthrough for identifying the transcriptome in non-model organisms [41]. If the research expenditure is sufficient, we recommend combining the long-read technology with scRNA-Seq reads for the identification and quantification of the novel transcriptome respectively.

### Conclusion

While most of the related studies focused on optimizing the quantification or assembly algorithms independently, few studies have discussed how the erroneous contigs generated by the assemblers affect the downstream analysis of RNA-Seq. In this study, we comprehensively examined the impact of biological complexity, assembly completeness and sequence ambiguity. We comparatively evaluated the performance of rnaSPAdes, Trans-ABySS and Trinity for *de novo* transcriptome assembly under three transcriptomes with different complexities. All of the selected assemblers showed a lower proportion of the fully-reconstructed transcripts as the complexity of transcriptome increases. In general, rnaSPAdes constructed the least number of contigs with the highest TransRate score, Trinity produced longer contigs, and Trans-ABySS generated the contigs with higher accuracy. As for quantification, we measured the reliability of RSEM, Kallisto and Salmon. The estimation made by three algorithms shows marginal differences. For each erroneous contig, the incomplete or over-extended contigs lead to unreliable estimation of the abundance of contigs. Moreover, we have found that if the redundant contigs are present in the assembly, the quantifiers tended to allocate the RNA-Seq reads to one of the duplicated contig. However, in rare cases, the quantifiers distributed the reads evenly to the contigs that share similar sequence content. On the contrary, the quantifiers tended to over-estimate the contigs that were assigned with multiple transcripts since the assemblers failed to distinguish the difference of these transcripts and reported only a single contig. To circumvent these issues, it is advisable to estimate the abundance on component-level rather than for individual transcripts. By exploring how these factors deteriorate the reliability of *de novo* RNA-Seq analysis, we provided valuable insights for the interplay between transcriptome assembly, quantification and sequence annotation. We anticipated these discoveries will be useful in the future development of assembly or quantification programs.

## Declarations

### Ethics approval and consent to participate

Not applicable.

### Consent for publication

Not applicable.

### Availability of data and material

The experimental datasets analyzed during the current study are available in the NCBI Short Read Archive repository, SRR453566 (https://www.ncbi.nlm.nih.gov/sra/SRR453566), SRR882109 (https://www.ncbi.nlm.nih.gov/sra/SRR882109), SRR203276 (https://www.ncbi.nlm.nih.gov/sra/SRR203276). The simulated datasets analyzed during the current study are available from the corresponding author on reasonable request.

### Authors’ contributions

P.-H.H. initiated the study, designed the analysis procedures, performed the analysis and wrote the manuscript. C.-Y.C. and Y.-J.O. commented on the draft and revised the manuscript. All authors read and approved the final manuscript.

## Acknowledgements

We wish to thank Yu-Chuan Chang and Mei-Ju May Chen for discussion on the analysis of quantification, Dr. Mong-Hsun Tsai for the comments on the analysis of RNA-Seq technology and Dr. Li-Yu Liu for the advices on statistical inference.

## Additional Files

**Additional File 1.** Title: Supplementary Tables. Description of data: Additional File 1: Table S1 to Table S6.

**Additional File 2.** Title: Supplementary Figures. Description of data: Additional File 2: Fig. S1 to Fig S6.

## Reference

Schena M, Shalon D, Davis RW, Brown PO: Quantitative monitoring of gene expression patterns with a complementary DNA microarray. Science 1995, 270(5235):467–470.

Mortazavi A, Williams BA, McCue K, Schaeffer L, Wold B: Mapping and quantifying mammalian transcriptomes by RNA-Seq. Nat Methods 2008, 5(7):621–628.

Wang Z, Gerstein M, Snyder M: RNA-Seq: a revolutionary tool for transcriptomics. Nat Rev Genet 2009, 10(1):57–63.

Genome KCoS: Genome 10K: a proposal to obtain whole-genome sequence for 10,000 vertebrate species. J Hered 2009, 100(6):659–674.

i KC: The i5K Initiative: advancing arthropod genomics for knowledge, human health, agriculture, and the environment. J Hered 2013, 104(5):595–600.

Zhang G, Rahbek C, Graves GR, Lei F, Jarvis ED, Gilbert MT: Genomics: Bird sequencing project takes off. Nature 2015, 522(7554):34.

Vijay N, Poelstra JW, Kunstner A, Wolf JB: Challenges and strategies in transcriptome assembly and differential gene expression quantification. A comprehensive in silico assessment of RNA-seq experiments. Mol Ecol 2013, 22(3):620–634.

Martin JA, Wang Z: Next-generation transcriptome assembly. Nat Rev Genet 2011, 12(10):671–682.

Conesa A, Madrigal P, Tarazona S, Gomez-Cabrero D, Cervera A, McPherson A, Szczesniak MW, Gaffney DJ, Elo LL, Zhang X et al: A survey of best practices for RNA-seq data analysis. Genome Biol 2016, 17:13.

Schulz MH, Zerbino DR, Vingron M, Birney E: Oases: robust de novo RNA-seq assembly across the dynamic range of expression levels. Bioinformatics 2012, 28(8):1086–1092.

Bankevich A, Nurk S, Antipov D, Gurevich AA, Dvorkin M, Kulikov AS, Lesin VM, Nikolenko SI, Pham S, Prjibelski AD et al: SPAdes: a new genome assembly algorithm and its applications to single-cell sequencing. J Comput Biol 2012, 19(5):455–477.

Xie Y, Wu G, Tang J, Luo R, Patterson J, Liu S, Huang W, He G, Gu S, Li S et al: SOAPdenovo-Trans: de novo transcriptome assembly with short RNA-Seq reads. Bioinformatics 2014, 30(12):1660–1666.

Robertson G, Schein J, Chiu R, Corbett R, Field M, Jackman SD, Mungall K, Lee S, Okada HM, Qian JQ et al: De novo assembly and analysis of RNA-seq data. Nat Methods 2010, 7(11):909–912.

Grabherr MG, Haas BJ, Yassour M, Levin JZ, Thompson DA, Amit I, Adiconis X, Fan L, Raychowdhury R, Zeng Q et al: Full-length transcriptome assembly from RNA-Seq data without a reference genome. Nat Biotechnol 2011, 29(7):644–652.

Papastamoulis P, Hensman J, Glaus P, Rattray M: Improved variational Bayes inference for transcript expression estimation. Stat Appl Genet Mol Biol 2014, 13(2):203–216.

Bray NL, Pimentel H, Melsted P, Pachter L: Near-optimal probabilistic RNA-seq quantification. Nat Biotechnol 2016, 34(5):525–527.

Li B, Dewey CN: RSEM: accurate transcript quantification from RNA-Seq data with or without a reference genome. BMC Bioinformatics 2011, 12:323.

Patro R, Duggal G, Love MI, Irizarry RA, Kingsford C: Salmon provides fast and bias-aware quantification of transcript expression. Nat Methods 2017, 14(4):417– 419.

Smith-Unna R, Boursnell C, Patro R, Hibberd JM, Kelly S: TransRate: reference-free quality assessment of de novo transcriptome assemblies. Genome Res 2016, 26(8):1134–1144.

Zhao QY, Wang Y, Kong YM, Luo D, Li X, Hao P: Optimizing de novo transcriptome assembly from short-read RNA-Seq data: a comparative study. BMC Bioinformatics 2011, 12 Suppl 14:S2.

Li B, Fillmore N, Bai Y, Collins M, Thomson JA, Stewart R, Dewey CN: Evaluation of de novo transcriptome assemblies from RNA-Seq data. Genome Biol 2014, 15(12):553.

Kanitz A, Gypas F, Gruber AJ, Gruber AR, Martin G, Zavolan M: Comparative assessment of methods for the computational inference of transcript isoform abundance from RNA-seq data. Genome Biol 2015, 16:150.

Zhang C, Zhang B, Lin LL, Zhao S: Evaluation and comparison of computational tools for RNA-seq isoform quantification. BMC Genomics 2017, 18(1):583.

Wang S, Gribskov M: Comprehensive evaluation of de novo transcriptome assembly programs and their effects on differential gene expression analysis. Bioinformatics 2017, 33(3):327–333.

Altschul SF, Madden TL, Schaffer AA, Zhang J, Zhang Z, Miller W, Lipman DJ: Gapped BLAST and PSI-BLAST: a new generation of protein database search programs. Nucleic Acids Res 1997, 25(17):3389–3402.

Nookaew I, Papini M, Pornputtapong N, Scalcinati G, Fagerberg L, Uhlen M, Nielsen J: A comprehensive comparison of RNA-Seq-based transcriptome analysis from reads to differential gene expression and cross-comparison with microarrays: a case study in Saccharomyces cerevisiae. Nucleic Acids Res 2012, 40(20):10084–10097.

Liu D, Xiong H, Ellis AE, Northrup NC, Rodriguez CO, Jr., O’Regan RM, Dalton S, Zhao S: Molecular homology and difference between spontaneous canine mammary cancer and human breast cancer. Cancer Res 2014, 74(18):5045–5056.

Griebel T, Zacher B, Ribeca P, Raineri E, Lacroix V, Guigo R, Sammeth M: Modelling and simulating generic RNA-Seq experiments with the flux simulator. Nucleic Acids Res 2012, 40(20):10073–10083.

Aken BL, Achuthan P, Akanni W, Amode MR, Bernsdorff F, Bhai J, Billis K, Carvalho-Silva D, Cummins C, Clapham P et al: Ensembl 2017. Nucleic Acids Res 2017, 45(D1):D635–D642.

Andrews S: FastQC: A quality control tool for high throughput sequence data. Reference Source 2010.

Bolger AM, Lohse M, Usadel B: Trimmomatic: a flexible trimmer for Illumina sequence data. Bioinformatics 2014, 30(15):2114–2120.

Li H, Durbin R: Fast and accurate short read alignment with Burrows-Wheeler transform. Bioinformatics 2009, 25(14):1754–1760.

Li B, Ruotti V, Stewart RM, Thomson JA, Dewey CN: RNA-Seq gene expression estimation with read mapping uncertainty. Bioinformatics 2010, 26(4):493–500.

Pachter L: Models for transcript quantification from RNA-Seq. arXiv preprint arXiv:11043889 2011.

Simpson JT, Wong K, Jackman SD, Schein JE, Jones SJ, Birol I: ABySS: a parallel assembler for short read sequence data. Genome Res 2009, 19(6):1117–1123.

Langmead B, Salzberg SL: Fast gapped-read alignment with Bowtie 2. Nat Methods 2012, 9(4):357–359.

Soneson C, Love MI, Robinson MD: Differential analyses for RNA-seq: transcript-level estimates improve gene-level inferences. F1000Res 2015, 4:1521.

Wang L, Si Y, Dedow LK, Shao Y, Liu P, Brutnell TP: A low-cost library construction protocol and data analysis pipeline for Illumina-based strand-specific multiplex RNA-seq. PLoS One 2011, 6(10):e26426.

Roberts A, Trapnell C, Donaghey J, Rinn JL, Pachter L: Improving RNA-Seq expression estimates by correcting for fragment bias. Genome Biol 2011, 12(3):R22.

Marinov GK, Williams BA, McCue K, Schroth GP, Gertz J, Myers RM, Wold BJ: From single-cell to cell-pool transcriptomes: stochasticity in gene expression and RNA splicing. Genome Res 2014, 24(3):496–510.

Rhoads A, Au KF: PacBio Sequencing and Its Applications. Genomics Proteomics Bioinformatics 2015, 13(5):278–289.

